# Genome-scale modeling of *Pseudomonas aeruginosa* PA14 unveils its broad metabolic capabilities and role of metabolism in drug potentiation

**DOI:** 10.1101/2021.04.15.439930

**Authors:** Sanjeev Dahal, Alina Renz, Andreas Dräger, Laurence Yang

## Abstract

*P. aeruginosa* is an opportunistic human pathogen that is one of the leading causes of hospital-acquired infections. We have developed an updated genome-scale model (GEM) of *Pseudomonas aeruginosa* PA14 for systems-study of the pathogen. We used both automated and semi-manual approaches to reconstruct and curate the model. After an extensive literature research, we added organism-specific reactions (e.g., phenazine transport and redox metabolism, cofactor metabolism, carnitine metabolism, oxalate production, etc.) to the model. This effort led to a highly curated, three-compartment, and mass-and-charge balanced BiGG model of PA14 that contains 1509 genes, 1779 metabolic reactions and 1151 unique metabolites. The model (*i*SD1509) has the largest genome coverage of *P. aeruginosa* PA14 to date with 424 more genes than the previous model (iPau1129). It is also the most accurate with prediction accuracies as high as 92.4% (for gene essentiality) and 93.5% (for substrate utilization). The model simulates growth in both aerobic and anaerobic conditions. It predicts the biosynthesis of the virulence factor phenazine as a process for the pathogen to grow in low-oxygen environment. Further, a mechanism for the overproduction of a drug susceptibility biomarker (gluconate) can be elucidated by the principles of optimal growth. Finally, the model also simulates drug activity potentiation and protection by fumarate and glyoxylate, respectively, and provides mechanistic explanations for these processes. Overall, *i*SD1509 can be utilized to decipher the metabolic mechanisms associated with virulence and antibiotic susceptibility of *P. aeruginosa* PA14 to aid in the development of effective intervention strategies.

## Introduction

*Pseudomonas aeruginosa* is a Gram-negative proteobacterium that is metabolically versatile and an opportunistic human pathogen. It is a leading cause of nosocomial infections [1, 2]. One of the well-known *P. aeruginosa* infection sites is in the lungs of cystic fibrosis (CF) patients. Such infections can lead to high morbidity and mortality [2]. The ability to resist multiple drugs (including aminoglycosides, quinolones and *β*-lactams), synthesize virulence factors (e.g., phenazines, proteases, lysins, exotoxins, etc.), and produce biofilms [1, 2, 3] allows *P. aeruginosa* to infect and colonize its host. The pathogenicity of *P. aeruginosa* and the host response can differ between strains [1].

*P. aeruginosa* PA14 is a hypervirulent strain and belongs to the most common clonal group [4]. A study in 2004 first published its genome, which is highly similar to *P. aeruginosa* PAO1’s genome, but carries two additional pathogenicity islands that contribute significantly to the virulence of PA14 [5, 6]. Therefore, the investigation of PA14 has gained particular interest in the research community.

Genome-scale metabolic models (GEMs) provide a reliable tool for systems study of bacteria [7, 8, 9]. GEMs can be advantageous in investigating pathogens because identifying potential intervention strategies can be challenging due to the wide range of genetic mutations and metabolic targets, and niche-specific alteration of metabolic processes [10, 11]. GEMs have been utilized for systems investigation of pathogenic species such as *Acinetobacter baumannii* [12, 13], *Klebsiella pneumoniae* [14], *Mycobacterium tuberculosis* [15], *Vibrio vulnificus* [16] and *P. aeruginosa* [17, 18]. For the development of GEMs, first a reconstruction of the metabolic pathways of the organism of interest is required, which can then be converted to a mathematical format that can be analyzed using constraint-based modeling and flux balance analysis approaches. A GEM of *Pseudomonas aeruginosa* PA14 (referred to as iPau1129) was published in 2017 in a study investigating the association between growth and virulence-linked pathways [17]. Several missing metabolic capabilities and standardization considerations in iPau1129 motivated us to reconstruct an updated model for PA14. For instance, iPau1129 does not possess multiple terminal oxidases, which are present in *P. aeruginosa* [19, 20, 21, 22]. Likewise, phenazine-dependent redox reactions including reactive oxygen species (ROS) formation were not included in iPau1129 [23, 24, 25]. We also sought to use human-interpretable metabolite and reaction identifiers standardized across a large repository of models—in particular, the Biochemical Genetic and Genomic (BiGG) standardization and modeling platform [26]. iPau1129 was constructed using ModelSEED identifiers without cross-referencing other databases.

In this study, we developed a reconstruction of PA14 strain using BiGG [26] identifiers with 424 additional unique genes compared with iPau1129. The model (*i*SD1509) demonstrated high prediction accuracy, and can also simulate growth in anaerobic environment. Furthermore, using the limed-FBA approach [27], we extended *i*SD1509 to investigate the effect of phenazine biosynthesis on the biomass production at various levels of oxygen availability. Finally, we also demonstrate the utility of *i*SD1509 in studying antibiotic resistance in *P. aeruginosa*.

## Materials and Methods

### Model simulations

For the simulation of the model, we performed flux balance analysis (FBA) on conditions reflecting the media used in a particular experimental study (SI Appendix, Table S1). The FBA simulations were performed using COBRApy package (v. 0.18.1) [28]. In most simulations, we used the aerobic biomass reaction as the objective function. For anaerobic growth predictions and comparison with aerobic growth rates, we changed the objective function to the anaerobic biomass reaction. If the biomass flux was computed to less than 10^−5^ h^-1^, “no growth” was assigned. For the computation of ubiquinone yield and growth rate comparison in aerobic and anaerobic environment, parsimonious FBA (pFBA) [29] was applied.

For the computation of gluconate production, both FBA and flux variability analysis (FVA) [30] (with loopless method and fraction of optimum set to 1) were applied. All simulations were performed in M9 medium. Experimental data to validate gluconate production were obtained from [31], where measurements for RpoN mutant and wild-type were performed in M9 media. Gluconate production for mutants besides RpoN were measured by [31] in SFCM media. However, for consistency and to facilitate interpretation, we simulated all mutants in M9 medium (a minimal medium) and found that these simulations were consistent with experiments in SFCM media (a rich medium).

To compare the wild-type and mutant fluxes, we applied a heuristic approach. First, any zero flux was replaced by 0.01. To compute gluconate yield, flux of gluconate secretion reaction was divided by that of glucose exchange reaction. For FBA simulations, the reaction fluxes or yields of the mutant were divided by those of the wildtype. For FVA fluxes, an average flux of each reaction was calculated by taking the mean of minimum and maximum flux values for both mutant and wildtype. Then, the mutant average reaction flux values were normalized by those of the wildtype.

### Superoxide leakage

We reconstructed the superoxide leakage by creating a net reaction composed of two: (1) normal reaction, in which water is produced as a by-product of redox metabolism (0.5 o2_e + pyoh2_e → h2o_e + pyo_e), and (2) leaky reaction in which superoxide is produced as a by-product (2.0 o2_e + pyoh2_e → 2.0 h_e + 2.0 o2s_e + pyo_e). We estimated the ROS leakage stoichiometric coefficient (*κ*) by taking into account the changes in the amount of pyocyanin (which can lead to superoxide production) and hydrogen peroxide (a product of superoxide dismutation) produced after 24 hours of growth [24]. According to this method,

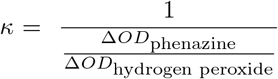

Then, we multiplied the stoichiometric coefficients of reaction (2) by *κ* and those of reaction (1) by 1-*κ* before adding up the reactions to generate a net reaction. The value of *κ* was estimated to be 0.29. Therefore, the net reaction used in the model is: 0.935 o2_e + pyoh2_e → 0.58 h_e + 0.71 h2o_e + 0.58 o2_e + pyo_e

### Addition of Unknown Reactions

For the addition of unknown reactions derived from the literature, we first checked for mass-and-charge balance. Then, we entered the reactions in the eQuilibrator program (v. 0.4.1) [32] in Python (v. 3.7.7). The parameters-pH = 7.5, ionic strength = 0.25 M, temperature = 25°C, control of magnesium ion (pMg) = 3 were used. Physiological concentrations (aqueous reactants at 1 mM) were assumed for the calculation of Gibbs free energy of transformation (ΔG^’m^). Reactions with ΔG^’m^<0 were added to the model.

## Results

### Model reconstruction and validation

The reconstruction of *i*SD1509 was developed using the pipeline outlined (SI appendix, Fig. S1 and Extended Results). We compared the reconstructions of *i*SD1509 and iPau1129. The *i*SD1509 model contains considerably higher number of genes, reactions, and metabolites than iPau1129 (Fig. 1*A*). It possesses 424 more unique genes than in iPau1129. When we compared the top KEGG pathways between the two reconstructions, the number of genes in almost every pathway was higher in *i*SD1509 than in iPau1129 (Fig. 1*B*). The current MEMOTE (v. 0.12.0) score of *i*SD1509 is 88% (SI Appendix, Fig. S2).

**Figure 1.**
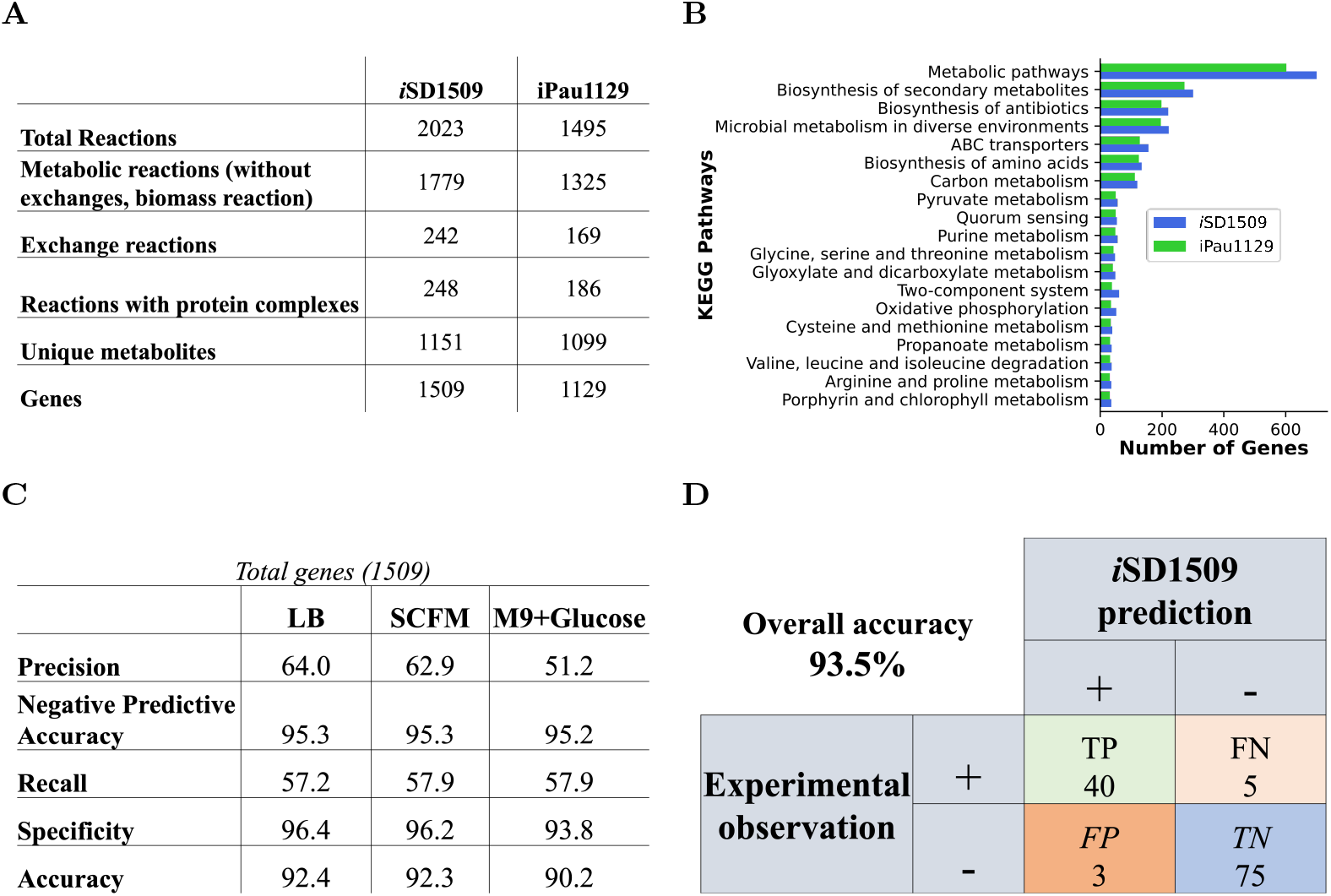
Genome-scale reconstruction of *Pseudomonas aeruginosa* PA14, *i*SD1509 and its predictive assessment. (*A*) When compared with iPau1129, the latest model *i*SD1509 has considerable increase in the reaction, gene, and metabolite content. (*B*) The top KEGG pathways between both reconstructions were compared, and *i*SD1509 evidently has higher gene content in almost every top KEGG pathway. (*C*) *i*SD1509 was then used for predicting core essential genes determined in a separate study [35]. The model prediction accuracy was found to be >90% in three different media conditions indicating a highly predictive model. (*D*) Using *i*SD1509, carbon substrate utilization was predicted for a new set of data [36]. The prediction accuracy was computed to be 93.5%.

Next, we validated *i*SD1509 by performing two tests: 1) substrate utilization in minimal media, and 2) gene essentiality. We used the same datasets utilized for validating iPau1129 [17].

The model can simulate growth on LB, SCFM (Synthetic Cystic Fibrosis Medium), and minimal media containing individual substrates (87 compounds). On the minimal medium, the model predicts substrate utilization with an accuracy of 94.3%. This is a significant improvement compared to the iPau1129 model (80.5%). The predicted growth rate on the glucose minimal medium (0.89 h^-1^) is within the range of experimentally determined growth rates for *P. aeruginosa* [33]. Next, we compared the metabolic flux analysis (MFA) results with the predicted FBA values for growth on glucose minimal medium. Due to the unavailability of MFA data for PA14 and since both PA14 and PAO1 strains are highly similar [5], we used the MFA data for PAO1 [34]. FBA predictions and MFA results were highly correlated (Pearson correlation coefficient: 0.91, *p*-value<0.001) (SI Appendix, Fig. S3).

For gene essentiality validation, we performed predictions in LB medium. For the genes common between *i*SD1509 and iPau1129 (1,085 genes), *i*SD1509 retained a similar accuracy (*i*SD1509: 90.5% vs. iPau1129: 91.0%) by achieving a >6% higher recall (SI Appendix, Fig. S4*A*). An increase in 1% accuracy was achieved if all the genes in *i*SD1509 were considered (SI Appendix, Fig. S4*B*).

### Predictive assessment of *i*SD1509

Next, we assessed the model using new datasets. For the gene essentiality data, 321 core essential genes identified in a recent transposon insertion sequence study were used [35]. Since the core essential genes were identified in three different media conditions– LB, SCFM and glucose minimal (M9) medium, we computed gene essentiality on all three conditions. For LB, SCFM, and M9 minimal medium, overall accuracies were 92.4%, 92.3%, and 90.2%, respectively. Precision ranged between 51.2% (M9) and 64.0% (LB), and recall ranged from 57.2% (LB) to 57.9% (M9 and SFCM) (Fig. 1*C*). Our model also demonstrated higher prediction accuracy than iPau1129 in LB medium for the common core essential genes (SI Appendix, Fig. S4*C*).

We assessed substrate utilization using the data published in Dunphy *et al*. [36]. We simulated the model for growth on 131 minimal media containing different substrates. On eight substrates, the two experimental data sets ([17] vs. [36]) disagreed on *P. aeruginosa*’s ability to grow. Hence, we excluded these eight substrates when assessing our model. On 123 substrates, the model simulated growth with a high accuracy of 93.5% (Fig. 1*D*).

### Anaerobic growth

*P. aeruginosa* can utilize nitrate or nitrite as terminal acceptors for growth in the absence of oxygen. When we first simulated the model in anaerobic condition, it could not simulate growth because the two biomass constituents-ubiquinone-9 (UQ9) and thiamine diphosphate (THMPP) could not be produced.

For anaerobic UQ9 production, a recent study [37] demonstrated that UQ9 can be produced by an alternate pathway within the UQ9 biosynthesis chain. The biosynthetic proteins are shared between aerobic and anaerobic pathways except for the ones that catalyze the three reactions associated with hydroxylation of intermediate metabolites (Fig. 2*A*). In an anaerobic environment, an alternative oxidizing molecule (proposed to be prephenate) could be involved [38]. Therefore, we added prephenate-based hypothetical reactions as anaerobic alternatives to the aforementioned three hydroxylation reactions. These reactions are mass-and-charge balanced, and their G^’m^s were computed using eQuilibrator (v. 0.4.1) [32] to determine their feasibility. For anaerobic THMPP production, no alternative pathway could be identified in *P. aeruginosa* by either literature search or annotation-based methods. Hence, we removed this metabolite and adjusted the biomass reaction creating an anaerobic biomass reaction.

**Figure 2.**
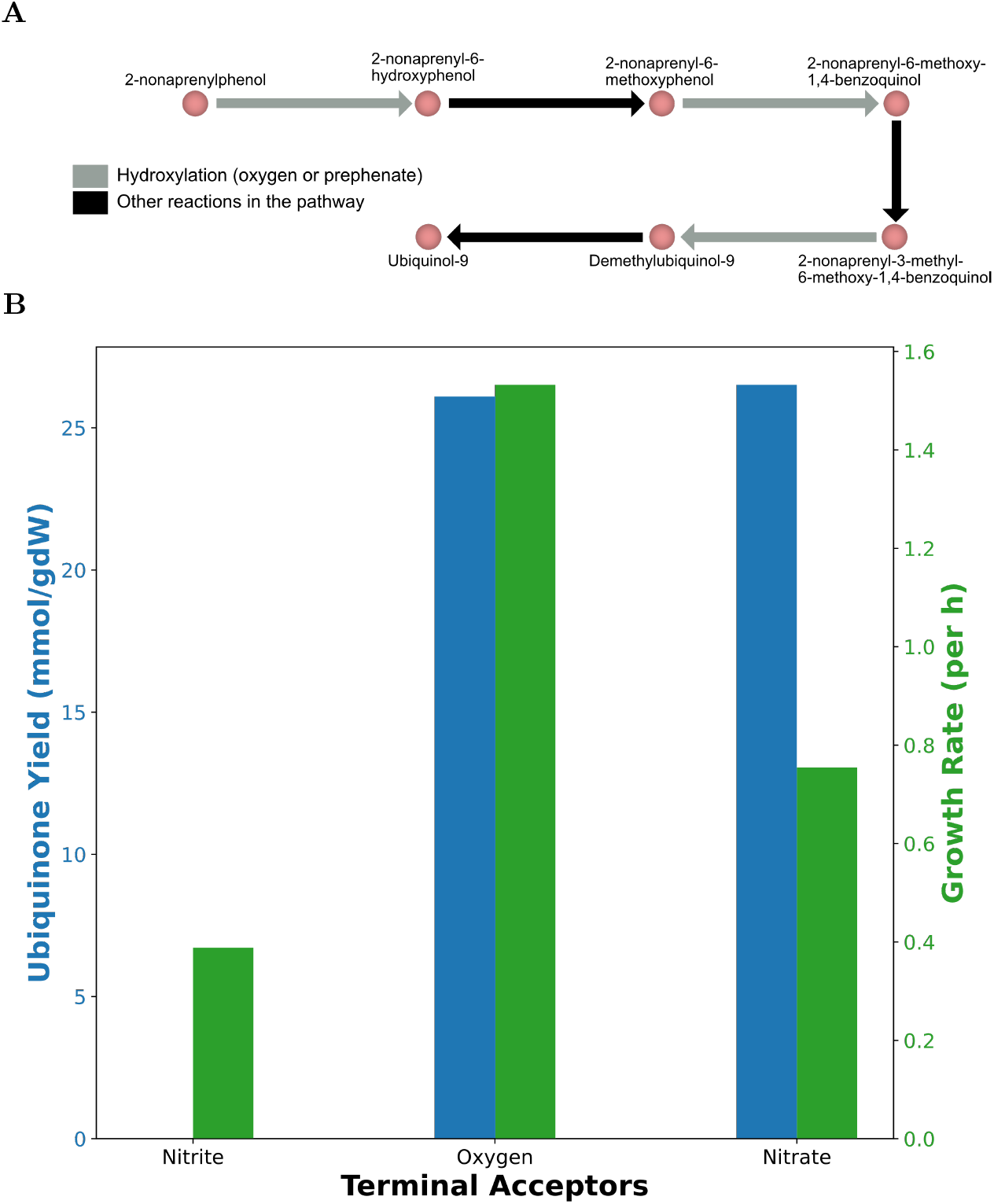
Our model predicts anaerobic growth of *P. aeruginosa* along with higher production of ubiquinone in anaerobic condition than in aerobic one. (*A*) The proposed pathway for ubiquinone production in both aerobic and anaerobic conditions share same enzymes except for the hydroxylation reactions [37, 38]. (*B*) The model simulates growth rates (green bars) in media with different terminal electron acceptors including nitrate and nitrite. Likewise, the model predicts that the ubiquinone (UQ9) yield (blue bars) in nitrate is slightly higher compared to that in aerobic conditions which agrees with the experimental data [37]. In nitrite, the UQ9 yield is extremely low (2.63 × 10^−4^ mmol gDW^-1^).

Following the changes, we could simulate anaerobic growth in LB medium. We then compared the growth rates between aerobic and anaerobic conditions by applying Pfba [29] and using an anaerobic biomass reaction as the objective function. We observed that the biomass production is lower under the anaerobic conditions than in the aerobic condition. (Fig. 2*B*). Studies have suggested that the energy yield of nitrate is lower than that of oxygen in *Pseudomonas* strains [39]. We also computed the UQ9 production in aerobic and anaerobic conditions. For this analysis, the reactions pertaining to ubiquinone-8 (UQ8) were turned off, and the flux-sum was computed for UQ9 which was then divided by the growth rate. The model predicted that the UQ9 production is slightly higher in the nitrate-supplemented medium than in the aerobic condition, which has been experimentally demonstrated (Fig. 2*B*)[37]. Our model also showed that the UQ9 yield in nitrite-supplemented medium is considerably low (2.63 × 10^−4^ mmol gDW^-1^), but the prediction has not been experimentally validated.

### Production of pyocyanin in *P. aeruginosa* can be simulated by extended *i*SD1509

We next used *i*SD1509 to investigate phenazine production in *P. aeruginosa* PA14. *Pseudomonas* can produce and secrete phenazines (e.g., pyocyanin), which possess redox properties and can cycle in and out of the cell. Usually, they are reduced within the cell, then they go to the extracellular space to get oxidized (by donating electrons to acceptors such as oxygen), and finally get transported back in the cell to complete a redox cycle [23, 40]. Studies have demonstrated that phenazine production is stimulated by low oxygen tension [41]. Further, phenazines are involved in the survival of *P. aeruginosa* in biofilm environment in which oxygen is limiting [23, 40].

To simulate the production of pyocyanin, which can act as a cofactor [42], we applied limed-FBA on *i*SD1509 to account for the dilution of this cofactor [27]. Since the cofactor is not part of the biomass reaction in *i*SD1509, it does not dilute as the model simulates growth. Hence, the model cannot predict the synthesis of pyocyanin as it is regenerated as part of the cofactor cycle (SI Appendix, Fig. S5*A*). Using *i*SD1509_limed, the oxygen-dependent synthesis of pyocyanin could be simulated (SI Appendix, Fig. S5*A*).

We utilized *i*SD1509_limed to study the effect of oxygen availability and phenazine production on the growth rate of *P. aeruginosa*. For the simulation, the oxygen import flux was constrained over a range (0.5 mmol gDW^-1^ h^-1^ to 10 mmol gDW^-1^ h^-1^). For each of those flux constraints, pyocyanin synthesis flux was also constrained over a range (0.000088 mmol gDW^-1^ h^-1^ (a tenth of the maximum value) to maximum value (0.00088 mmol gDW^-1^ h^-1^ computed for 0.5 mmol gDW^-1^ h^-1^ oxygen import flux)). For each value of oxygen import flux and pyocyanin biosynthesis flux, the growth rate was computed. With this analysis, we observed that for lower oxygen import flux (less oxygen available to the cell), the effect of pyocyanin production on the growth rate is pronounced. In contrast, when more oxygen is available to the cell, pyocyanin synthesis does not considerably contribute to the biomass production (Fig. 3 and SI Appendix, Fig. S5*B*).

**Figure 3.**
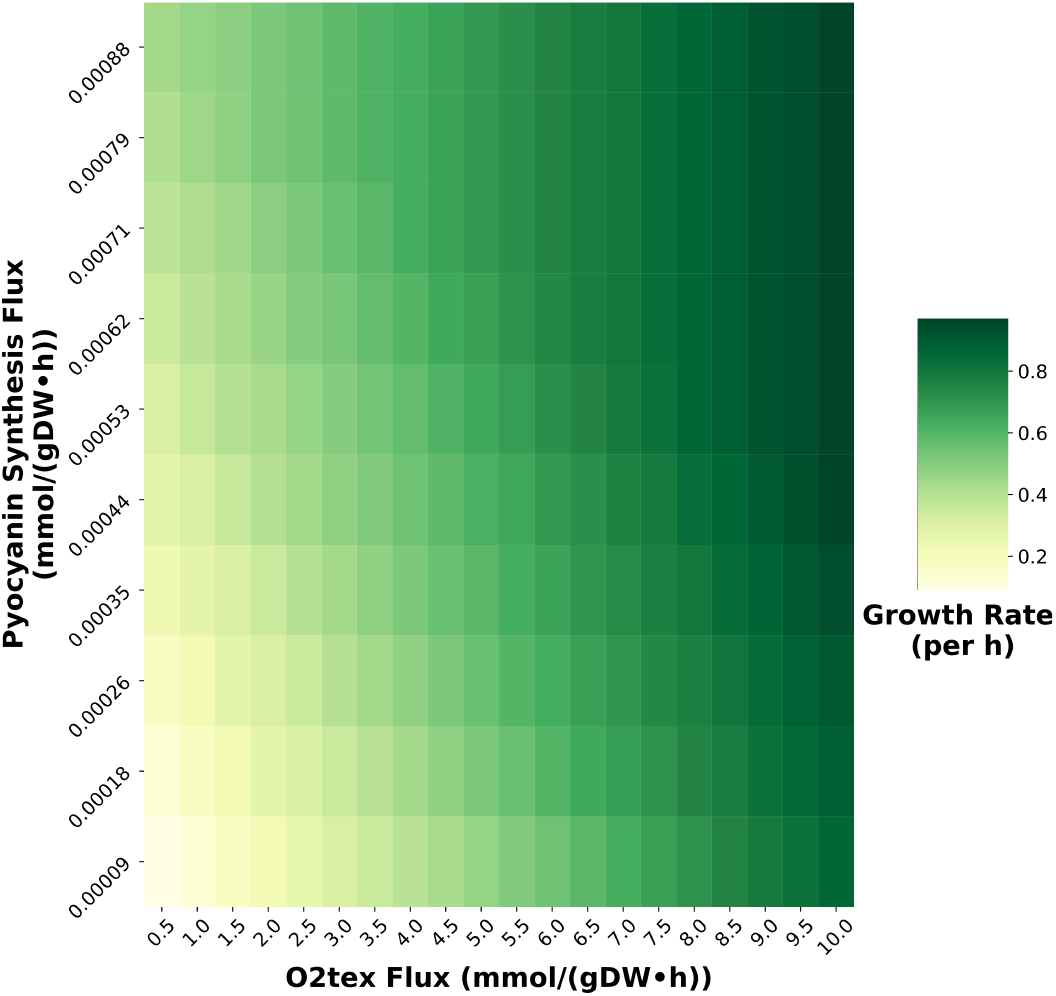
The limed-FBA model can predict the biomass production dependence on pyocyanin biosynthesis and oxygen availability. Simulations were performed over a wide range of oxygen uptake (from extracellular to periplasm) and pyocyanin biosynthesis flux constraints by optimizing for biomass production. For oxygen-limited condition (lower oxygen import flux), the effect of pyocyanin production on the growth rate is more profound compared to that in more oxygen-rich condition (higher oxygen import flux).

The *i*SD1509_limed can mechanistically explain the production of pyocyanin at low oxygen availability. In such environments, *Pseudomonas* is forced to divert resources towards phenazine production to counteract the imbalance in the intracellular redox state caused by the reduced availability of oxygen, which has been demonstrated experimentally [23]. Therefore, phenazines are crucial to the survival of the pathogen at low oxygen availability (e.g., within biofilms). We can, hence, use *i*SD1509_limed to identify potential intervention strategies that target phenazine redox cycle (SI Appendix, Fig. S5*C*).

### Identification of potential mutants that overproduce gluconate

The *rpoN* mutant is a significant gluconate producer. The loss of function mutation in *rpoN* is common among the clinical isolates in CF patients [43, 44]. Behrends *et al*. proposed that gluconate production is positively (weak but significant) correlated to reduced antibiotic susceptibility. Further, the investigators demonstrated that only one indirect target of RpoN, 6-phosphogluconate dehydratase (6PGDH, PA14_22910) could replicate the gluconate overproduction phenotype of the *rpoN* mutant [31].

We used *i*SD1509 to recapitulate the results of the study [31] on the glucose minimal (M9) medium. Of the knockouts of the thirteen indirect targets of RpoN, the FBA simulations accurately predicted that only the deletion of 6PGDH (*edd* gene) leads to a significant increase in the gluconate production compared to the wildtype (Fig. 4*A*). The model also correctly showed that the growth of the *edd* mutant is considerably affected. The simulations provided insights into other mutants such as deletion of gluconate symporter gene (PA14_34630) leading to increased flux through glucose transport reaction (GLCabcpp) (Fig. 4*A*).

**Figure 4.**
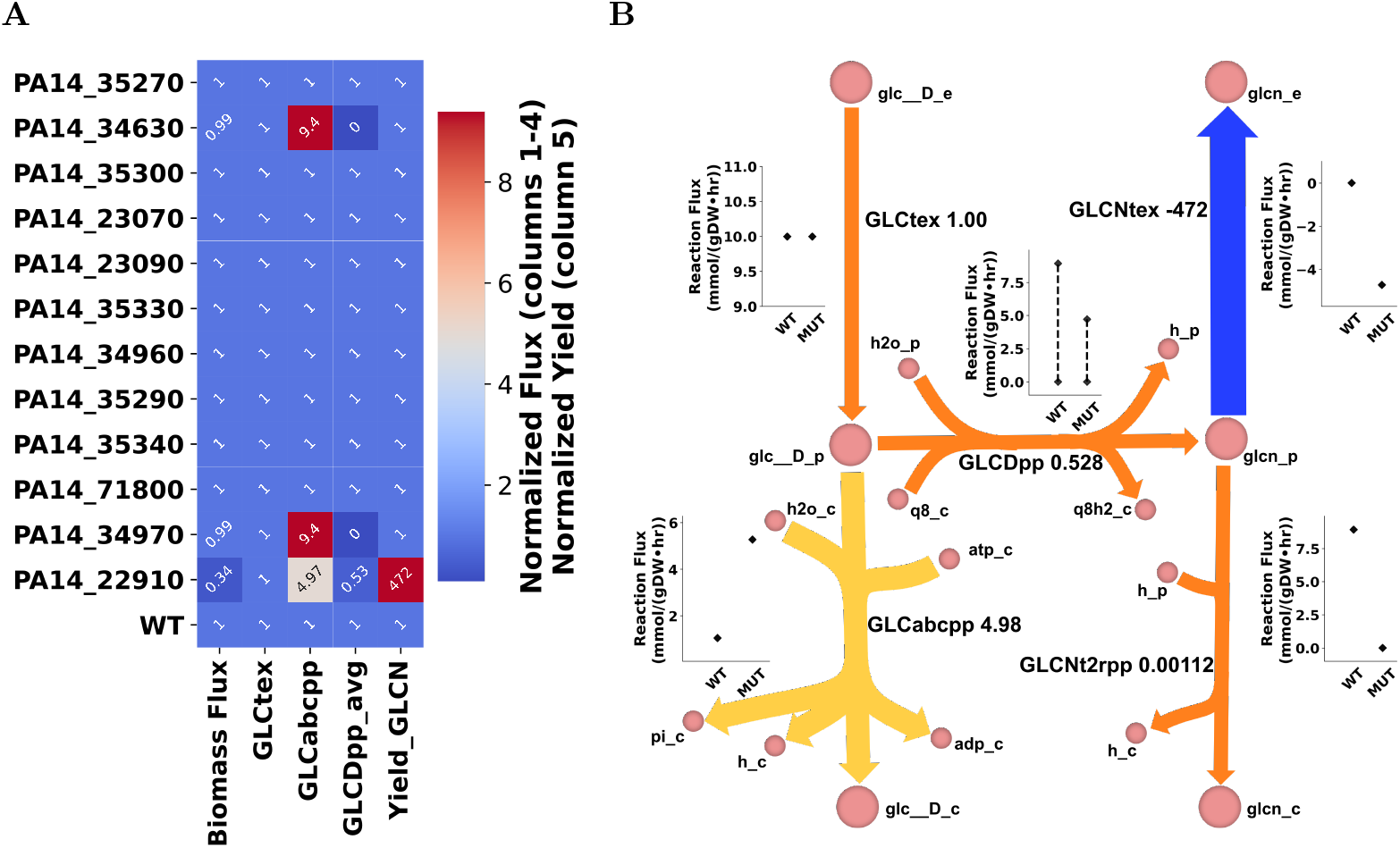
Our model can accurately predict gluconate production in the mutants of the genes regulated by RpoN. Thirteen genes regulated by RpoN were *in silico* knocked out in this study in order to recapitulate the results of experimental study [31]. (*A*) FBA simulations were able to accurately predict that only *edd* mutant (PA14_22910) produces considerable amount of gluconate compared to the wildtype. Furthermore, the decrease in the growth rate of the mutant was also recapitulated in this study. For calculations, zero flux values were first replaced with 0.01. The fluxes of the reactions catalyzed by glucose dehydrogenase (GLCDpp and GLCDpp q9) have been averaged (column GLCDpp avg).For gluconate yield (mmol gluconate per mmol glucose, last column), gluconate secretion flux was divided by glucose exchange flux. Then, mutant reaction fluxes/yield were divided by respective wildtype reaction fluxes/yield. (*B*) To further characterize the *edd* mutant, FVA simulations were carried out in order to examine the flux range of desired reactions in both wildtype and mutant. Gluconate excretion is required by the mutant for optimal growth in the given condition. Furthermore, the glucose flux in the mutant is divided between glucose transport and glucose dehydrogenase reactions whereas in the wildtype, majority of the flux is channeled towards the Entner-Doudoroff pathway through glucose dehydrogenase reaction (SI appendix, Fig. S4). The values (after the reaction names) are derived by dividing the average reaction fluxes of the *edd* mutant by those of the wildtype. To avoid division by zeroes, all the zeroes were replaced by 0.01.

Next, we performed FVA simulations to determine whether gluconate production in *edd* mutant was a requirement for the optimal growth. The gluconate secretion (GLCNtex) reaction flux range (absolute value) in the mutant (4.72 mmol gDW^-1^ h^-1^) is fixed at a higher rate than in the wildtype (0 mmol gDW^-1^ h^-1^) suggesting that for the optimal growth of the mutant, gluconate is forced to be transported out of the cell.

### Glyoxylate shunt decreases the TCA flux leading to lower respiration rate and lower proton motive force

Meylan *et al*. [45] showed that the addition of fumarate along with tobramycin leads to greater drug uptake and activity whereas glyoxylate protects the cells from tobramycin lethality. The study showed that the higher tobramycin uptake and activity in fumarate-containing medium is due to greater proton motive force (PMF) and increased respiration rate caused by higher flux through the TCA cycle, respectively. Likewise, in glyoxylate-containing medium, glyoxylate, by directly inhibiting *α*-ketoglutarate dehydrogenase, diverts the flux away from TCA cycle towards the glyoxylate shunt leading to reduced TCA cycle activity. We used *i*SD1509 to test an alternative, i.e., whether the law of optimal growth can explain drug protection by glyoxylate supplementation. Furthermore, we used *i*SD1509 to study the pathway utilization differences between the two metabolite supplementations.

We simulated *i*SD1509 on minimal (M9) medium containing glyoxylate or fumarate, and low amounts of citrate by optimizing for the biomass production. Since glyoxylate was not predicted to be a growth-inducing substrate in the substrate utilization assessment step, an artificial glyoxylate uptake reaction was added. We performed a flux sampling analysis by constraining the biomass flux to 90% of the estimated growth rate (by FBA) on both media. For proton flux calculations, we computed the flux-sum of periplasmic proton. Then, we compared the median fluxes of the reactions pertaining to TCA cycle and glyoxylate cycle between the two media conditions. The model predictions agreed with Meylan *et al*. [45] that the glyoxylate shunt indeed drives the flux away from TCA cycle in glyoxylate-containing medium. Likewise, the simulations also indicate that the glyoxylate flux is diverted towards the reactions catalyzed by malic enzymes, which recycle the necessary cofactors— NADH and NADPH. Unlike the Meylan study, the model predicted higher flux activity through pyruvate dehydrogenase reaction in the glyoxylate minimal medium. Moreover, the production of oxalate and glycolate were not confirmed by the model predictions. Instead, the glyoxylate flux was diverted towards the reaction catalyzed by glyoxylate carboligase. We also observed that the oxygen uptake rate and proton flux-sum were higher in the fumarate-than in the glyoxylate-supplemented medium, which led to increased drug uptake and activity in fumarate treatment (Fig. 5). Therefore, the law of optimal growth could also explain drug potentiation and drug protection by fumarate and glyoxylate supplementation, respectively.

**Figure 5.**
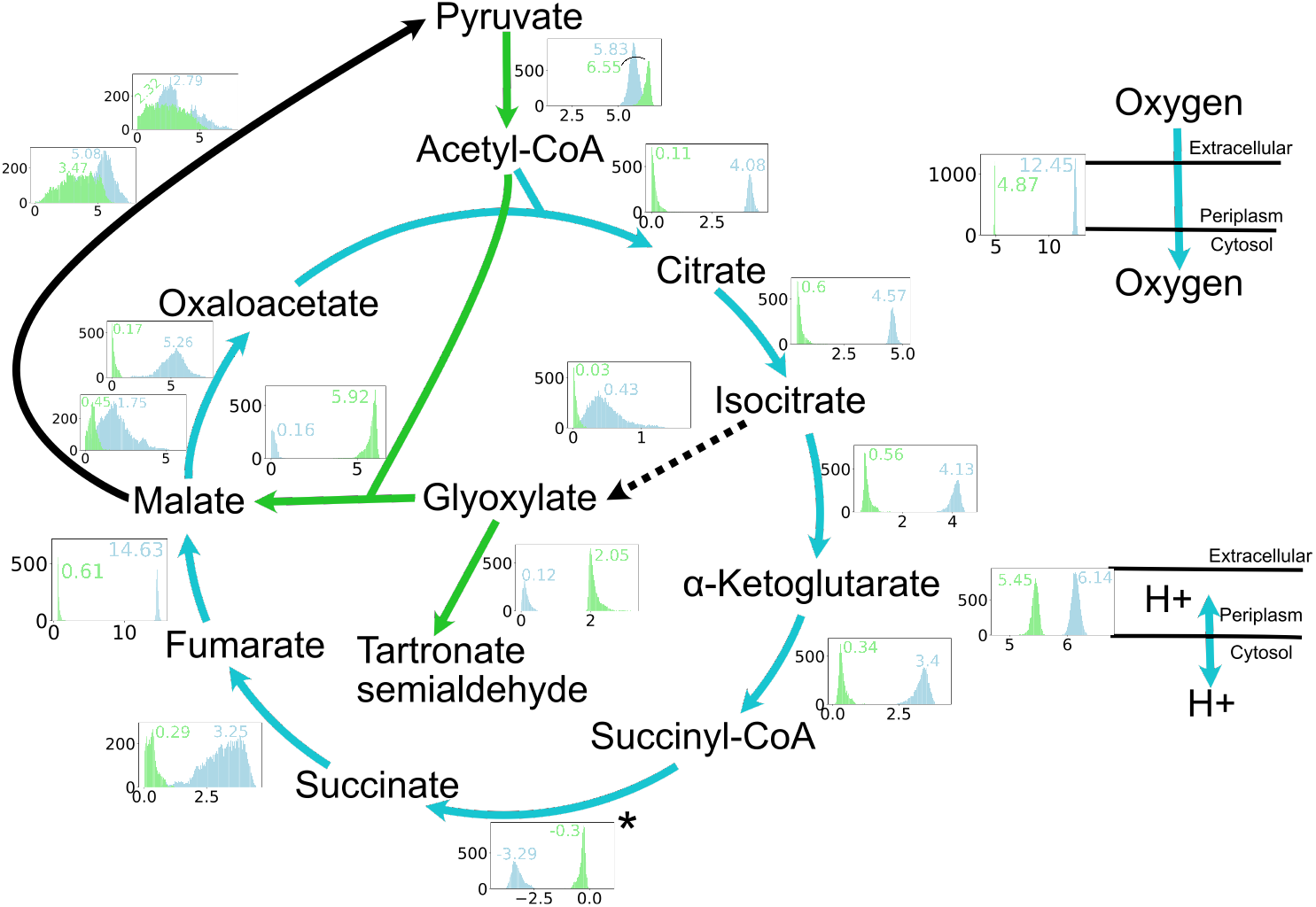
The *i*SD1509 model predicts that the glyoxylate shunt causes lower flux through the TCA cycle leading to lower oxygen uptake rate and PMF. Flux sampling analysis was performed in minimal media containing fumarate and glyoxylate to identify the flux distributions of TCA cycle reactions. According to the analysis, the TCA cycle is significantly upregulated in fumarate-supplemented medium compared to the glyoxylate one. The asterisk for reversible succinyl-CoA synthetase reaction is to note that the reaction flux is negative. In glyoxylate, the flux from acetyl-CoA is shunted towards the glyoxylate cycle as shown in the figure. The flux goes through the reactions catalyzed by malic enzymes to regenerate the cofactors— NADH and NADPH. This leads to a reduced TCA flux, decreased oxygen uptake rate, and PMF. All the sampling results (boxes with flux distributions where green: glyoxylate and blue: fumarate) are for reaction fluxes except for proton for which flux-sum yield was computed from the sampling data. Arrow colors indicate median fluxes being higher in one condition versus the other such that green: glyoxylate, blue: fumarate, black and bold: similar, and black and dotted: low flux in both conditions.

## Conclusion

*Pseudomonas aeruginosa* is an extensively studied organism in association with human infections especially in CF lungs. Since it is known to possess multiple mechanisms of drug resistance (e.g., lipopolysaccharide modification, overexpression of efflux pumps) [3] and an ability to survive within the biofilm environment, eradication of *Pseudomonas* infections can be challenging. Even though *P. aeruginosa* PAO1, a laboratory strain, has been widely investigated, and a copious amount of data is available for the organism, such progress has not been made for other *P. aeruginosa* organisms including the hypervirulent strain PA14. Using a predictive model such as *i*SD1509, which contains organism-specific knowledge, we can not only bridge the gaps but can also provide a platform to design experiments and strategies to combat infections caused by PA14 strain.

In this study, we have demonstrated that *i*SD1509 is a highly predictive and reliable model with accuracies of 90.2-93.4% and 93.5% for gene essentiality and substrate utilization predictions, respectively. Further, guided by the model simulations, we predicted that the biosynthesis of ubiquinone-9 is required for the anaerobic growth of PA14, and a recent study [37] corroborated this hypothesis. The model also predicted that the survival of *P. aeruginosa* in low oxygen environment (such as within biofilms) is possible due to phenazine production and provided mechanistic insight into their biosynthesis.

The model successfully identified the mutant that produced higher amounts of gluconate than the wildtype among thirteen candidates. Using *i*SD1509, more computational experiments can be designed to identify other mutants that overproduce gluconate. Likewise, other possible biomarkers of drug resistance can be predicted. Furthermore, using multi-strain modeling approaches [46], models of various *P. aeruginosa* isolates can be simulated to compare biomarker production simultaneously.

Finally, we used *i*SD1509 to recapitulate the potentiation of an antibiotic by metabolite supplementation [45]. The model correctly differentiated between metabolites that increased drug lethality versus those that did not and offered mechanistic explanations for these responses. Namely, supplementing glyoxylate diverted flux away from the TCA cycle, lowering the respiration rate that consequently led to lower drug activity. Supplementing the medium with fumarate caused higher oxygen uptake rate due to higher TCA cycle activity which leads to higher drug activity [45]. Likewise, PMF (and ultimately drug uptake) was higher in the fumarate-containing medium than in the glyoxylate-containing medium. These results suggest that our model can be used to design antimicrobial strategies based on metabolic mechanisms, including metabolite supplementation, against *P. aeruginosa*.

In conclusion, this model can be seen as a computational platform to design experiments targeting *P. aeruginosa* metabolism at various growth states including within the biofilm environment. Likewise, novel computational targets of *P. aeruginosa* (i.e., pyocyanin redox cycle) can now be investigated to identify the best strategy to inhibit the growth of the pathogen. Overall, we expect the model to significantly accelerate our understanding of *P. aeruginosa* to combat the associated infections.

## Supporting Information Appendix (SI)

The materials and methods are detailed in SI Appendix, including reconstruction of *i*SD1509, simulations using the model, which were performed in COBRApy package (v. 0.18.1) [28].

## Data availability

The model and sample codes are available at https://github.com/dahalsanzeev/iSD1509-M. All other study data are included in the article and/or supporting information.

## Acknowledgments

This work was supported by Queen’s University and the Natural Sciences and Engineering Research Council of Canada (NSERC) [RGPIN-2020-06325]. AR and AD received support from the German Center for Infection Research (grant number 8020708703) and the Cluster of Excellence CMFI (Controlling Microbes to Fight Infections), project number EXC-2124/05.037_0, funded by the Deutsche Forschungsgemeinschaft (DFG, German Research Foundation) under Germany’s Excellence Strategy – EXC 20124 – 390838134.

## Extended Results

### Initial reconstruction and manual refinement

We used CarveMe [47] to create a draft reconstruction that contained 2196 reactions and 1482 genes. Following this, we added the biomass reaction from the previous SEED model (iPau1129) [17]. We gap-filled our model in LB rich medium to produce all the biomass constituents by adding the reactions from iPau1129 or from other BiGG models.

Next, we curated our model through a semi-automated approach. We chose seventeen different BiGG models to perform a bidirectional blast hit for each of the reactions present in the model (Table S2). We then manually checked the reactions for any discrepancy in the gene-protein-reaction (GPR) associations made by CarveMe and by our approach. Any discrepancy was resolved using the knowledge from other databases (e.g., IMG, KEGG). Reactions removed from the model are provided (Dataset S1).

Using carbon substrate essentiality [17] and gene essentiality data [48], we iteratively filled more knowledge gaps either through annotation- or literature-derived information. A list of reactions that were added to the model is provided (Dataset S2). We also utilized gene essentiality data to fix GPR associations. Likewise, biomass reaction was also modified during this process to better reflect the gene essentiality data.

We next added any extra genes in iPau1129 not present in our model. This process led to the addition of reactions related to alginate production, rhamnolipid production, pyochelin production, etc. Only 44 genes from iPau1129 are missing in *i*SD1509 (Dataset S3). Finally, we checked and corrected any mass-and-charge imbalance, first by applying the knowledge from well-curated BiGG models (*i*JN1462 [49] and *i*ML1515 [50]), which lead to only 129 metabolites that required additional manual curation. The formula and charges of those metabolites were changed either using information from metabolite databases (BiGG, MetaNetX, PubChem or ModelSeed) or manually by mass-and-charge balancing the reactions (Dataset S4).

### Literature derived curation and addition of reactions

We curated the knowledge derived from literature to add to the reconstruction to make the model more strain-specific. The reactions pertaining to multiple terminal oxidases, reactive oxygen species elimination, n-alkane degradation, phenazine-associated metabolism, oxalate production, rubredoxin-based metabolism and H+-translocating NADH:quinone oxidoreductases were added. This led to the development of a model possessing 1509 genes. The model (referred to as *i*SD1509) contains 2023 total reactions and 1151 unique metabolites and possesses three compartments.

We analyzed the additional genes in *i*SD1509 by enrichment in both KEGG and COG categories (Fig. S6). In KEGG, notable enriched pathways were oxidative phosphorylation, sulfur metabolism, folate biosynthesis, glyoxylate and dicarboxylate metabolism, and biofilm formation. In COG, the top three categories were energy production and conversion (C), amino acid transport and metabolism (E), and inorganic ion transport and metabolism (P). We also examined the top 20 KEGG pathways of the new metabolic reconstruction (Fig. S6*C*). The greatest number of genes in *i*SD1509 belong to the category “Metabolic pathways.” Interestingly, other significant pathways are associated with biosynthesis of secondary metabolites, metabolism in diverse environments, biosynthesis of antibiotics, indicating a reconstruction of a metabolically versatile organism (Fig. S6*C*).

## Extended Materials and Methods

### Initial Reconstruction

For the initial reconstruction, CarveMe [47] with gap-filling function in LB medium was utilized. From the previous *P. aeruginosa* model (iPau1129) [17], we added the biomass reaction along with other required reactions to simulate the growth of the model on LB medium. For this process, we first converted the ModelSEED [51] metabolite identifiers of iPau1129 to BiGG identifiers. Furthermore, the bounds of non-growth associated reaction were also added from iPau1129. Then, we removed all the artificial sink and demand reactions added by CarveMe by making sure that the respective metabolites can be produced or consumed by added reactions which are supported by gene evidence.

Then, the *Pseudomonas aeruginosa* PA14 reactome was inspected for the gene-protein-reaction (GPR) associations using a custom pipeline (Fig. S1). First, seventeen models from BiGG database and their respective protein FASTA files were downloaded (Table S2). Then, we performed a bidirectional blast hit (BBH) for all the reactions present in the model by prioritizing the strains that are taxonomically closer to PA14 strain. We applied a stringent method such that only the top hits in both directions were considered as the correct gene. If not, we manually insepcted the top blast hits in different databases including IMG [52] and KEGG [53]. These GPR associations were then used as alternatives to CarveMe predictions. We performed a rigorous manual check for the reactions whose GPR associations were derived from the BBH approach from models other than those from *Pseudomonas putida*. For any discrepancy between GPR associations from CarveMe and those from BBH approach, we checked IMG and/or KEGG and/or iPau1129 model to assign correct gene associations. Any reactions that did not have associated GPRs or with no evidence to be present in PA14 were discarded in this process. We also removed any unnecessary loops during this process. The list of deleted reactions is provided (Dataset S1). We iteratively improved and validated the model using substrate utilization [17] and gene essentiality [48] data which led to the modification of the biomass reaction and more changes to GPR associations. We also added any genes from iPau1129 that were not associated with the model at that time.

### Reaction Mass and Charge Balance

For mass-and-charge balance, metabolite formula and charge were assigned using *i*JN1462 first, and then *i*ML1515. Next, we manually checked the remaining metabolites on multiple databases including BiGG, MetaNetX, PubChem and ModelSEED to identify the correct formula and charge. Finally, if the information for metabolites could not be found in the aforementioned databases, we assigned the formula and charge by balancing the reactions that contain only those metabolites as the sole undetermined ones.

### Manual Reconstruction

Reactions related to anaerobic metabolism, phenazine-associated metabolism, terminal oxidases and alternative terminal oxidases, thiamine metabolism, nucleotide metabolism, n-alkane metabolism, oxalate production, anaerobic quinone production, rubredoxin-based metabolism, reactive oxygen species (ROS), and H+-translocating NADH:quinone oxidoreductases were added using annotation in KEGG and/or extensive knowledge from literature. All the added reactions are listed (Dataset S2). This reconstruction effort led to the development of the model, *i*SD1509.

### Substrate Utilization Screening and Gene Knockout Study

For the initial carbon source catabolic activity and gene essentiality validation, the same datasets used in Bartell *et al*. [17] were utilized. For substrate utilization, we simulated *i*SD1509 in minimal medium (Table S1) containing individual substrates and optimized for maximum biomass production. Since ModelSEED identifiers were used in Bartell *et al*., some of the substrate identifiers could not be converted to the BiGG ones, and hence were removed from this analysis. Overall, 87 compounds were compared. The media composition was provided generously by Papin lab. For gene essentiality comparison, we simulated the model in LB rich medium.

For substrate utilization screening and gene essentiality assessment of *i*SD1509, data collected from separate experimental studies were applied. For carbon source assay, the dataset by Dunphy *et al*. [36] containing 190 carbon sources was collected. In the collected dataset, since only general metabolite names were provided, the metabolites whose BiGG identifiers could be determined with high confidence were utilized for prediction. Furthermore, since the two experimental datasets ([17] and [36]) showed discrepancy in *P. aeruginosa*’s ability to grow on eight substrates, those compounds were removed from the analysis. Hence, the model was simulated on minimal medium containing 123 individual substrates for comparison. For assessing the gene essentiality, we used the dataset from a recent study by Poulsen *et al*. [35]. In this dataset, core essential genes were defined as essential genes in five media conditions and nine different strains. We analyzed these core essential genes in LB medium, SCFM (Synthetic Cystic Fibrosis Medium) and glucose minimal medium. For gene essentiality comparison in glucose minimal medium, iron had to be added even though the media used in the study [35] presumably did not contain iron.

The metrics for the comparison of model predictions and experimental data are defined as follows:

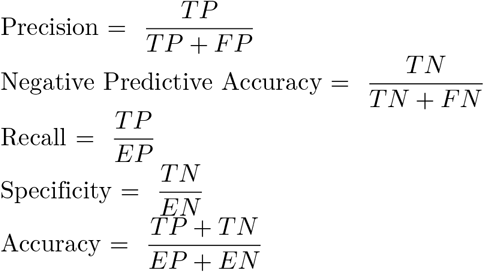

where, TP: true positive, FN: false negative, FP: false positive, TN: true negative, EP: experimental essential genes, and EN: experimental non-essential genes.

### Flux-sum Analysis

For the metabolites of interest, we performed flux-sum analysis [54]. Briefly, flux-sum (Φ_*i*_) for metabolite i can be computed using the following formula,

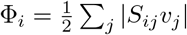

where j is the reaction in which the metabolite i participates in.

For the computation of yield, the flux-sum was either divided by the growth rate (for quinone production) or by the substrate flux multiplied by the number of carbon atoms in the substrate (for proton motive force (PMF)).

### Flux Sampling

We applied flux sampling approach [55] in the COBRApy package [28] by using optGpSampler [56] with 10000 samplings in both fumarate and glyoxylate minimal media. The growth rate was constrained to 90% of the rate predicted in FBA simulations in the respective media condition. By using the validate method of the optGpSampler, only valid samples were kept for further analysis. Furthermore, we used the autocorrelation plots and trace plotting for convergence analysis. To compare the distributions of reaction fluxes or metabolite flux-sums between the two different media conditions, we performed rank-sum tests.

### limed-FBA

For the simulation of dilution of cofactors, we used limed-FBA approach [27]. Briefly, all the reactions were first made irreversible. Then, for the reactions consuming or producing the desired cytosolic cofactor, a small dilution constant (*ϵ*) was applied such that

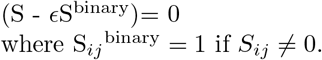

and *ϵ*_*ii*_ for metabolite *i*was computed using the following formula,

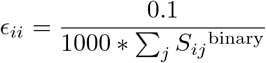

To not double-dilute biomass related metabolites, their *ϵ*_*ii*_ was made zero. Likewise, for simple transport reactions in which no chemical transformations (e.g., glucose(extracellular) → glucose (periplasm)) occur, their binary coefficients were assigned zeros to avoid any high-flux loops.

### Visualization of flux data

We first created a preliminary Escher [57] map in the Python-based framework, and then exported the map to web-based Escher tool for appropriate changes. Default settings were applied except for the customization of color and size of the edges for better visual representation. Then, we modified the Escher maps using vector graphic tool (Affinity Designer, v. 1.9.3).

We produced the heatmaps using *seaborn* package (v. 0.10.1) in Python. All the respective flux or yield values were divided by their respective wild-type values. To avoid division by zeroes, all the zero fluxes were first replaced by 0.01. For color range, robust quantile computation function was applied.

**Figure S1.**
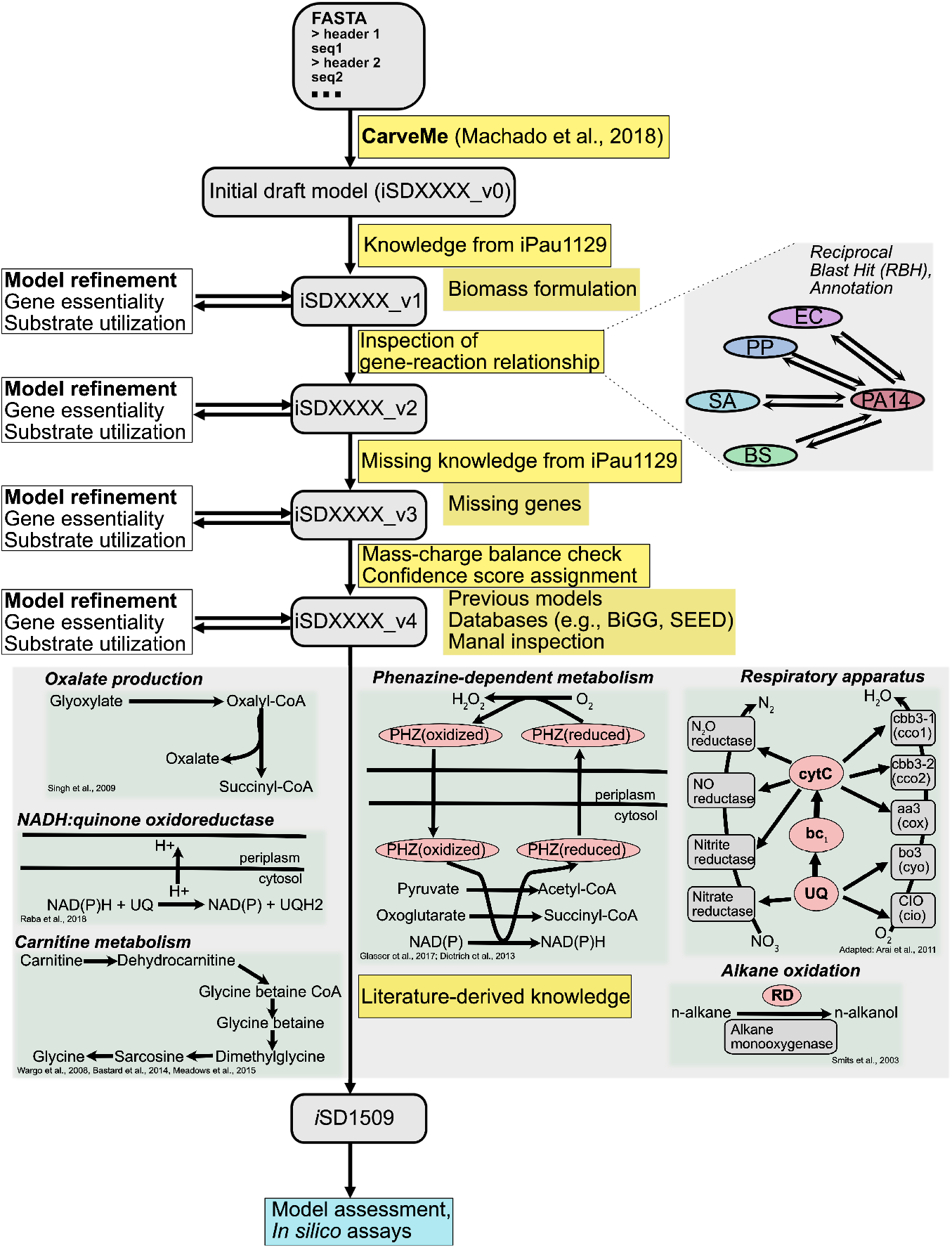
The working pipeline for building the genome-scale reconstruction of *P. aeruginosa* PA14. Both automated and semi-automated methods were applied in this pipeline, and the reconstruction was gap-filled using iPau1129 model wherever necessary. Furthermore, strain-specific reactions were added after extensive literature curation. At each stage, the model was validated and improved using gene essentiality and substrate utilization data from [17].

**Figure S2.**
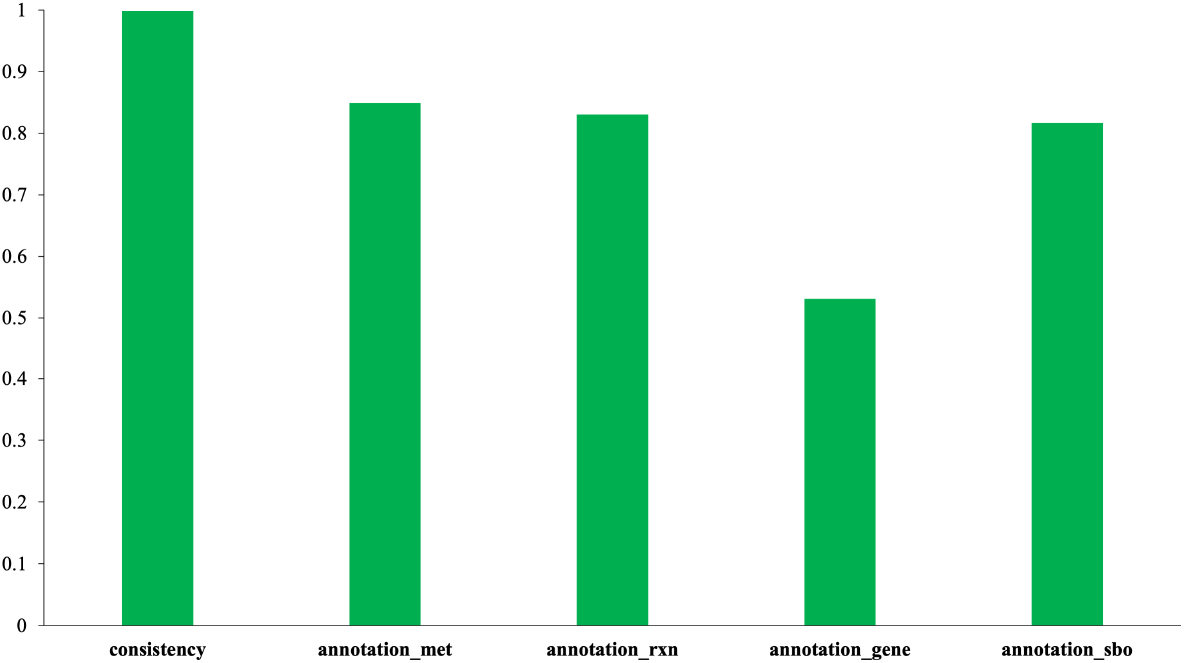
Memote scores for *i*SD1509. Overall, the score for the model is 88%.

**Figure S3.**
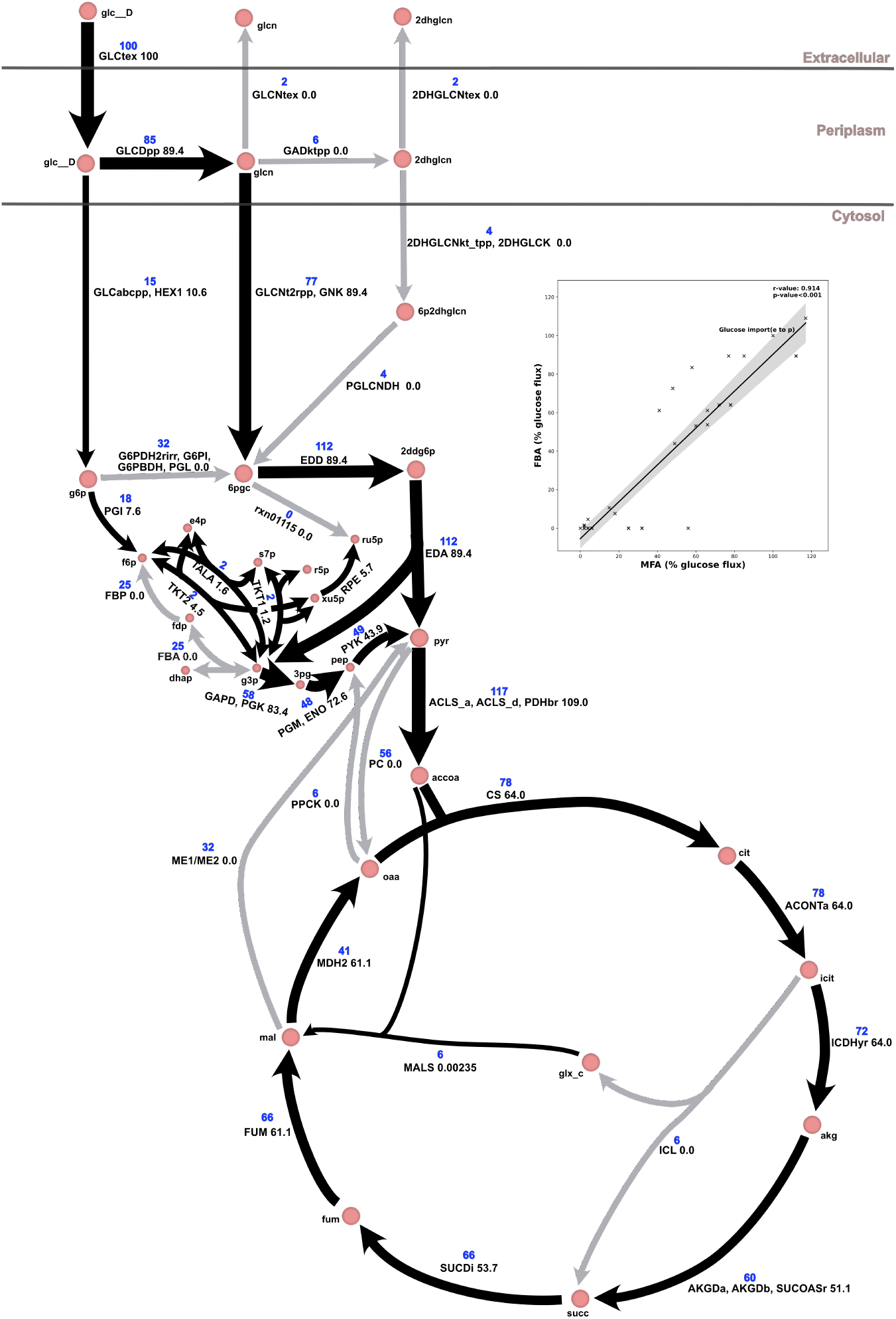
FBA simulations agree with MFA experimental data [34]. In the figure, the width of arrows represent the percentage of glucose flux. Only absolute values are shown here. All the zero value arrows are made grey for clarity. The MFA average values are written in the blue font if their respective reactions were found in the reconstruction. Lumped reactions are represented by multiple reactions separated by commas. Since two possible reactions (ME1 and ME2) are present in the M-model for the reactions catalyzed by the malic enzyme, both are shown separated by “/”. The box inset is a correlation plot that corroborates the fact that the majority of the predictions agree with the MFA results (correlation coefficient of 0.91 (p<0.001)). Grey bands denote 95% confidence interval. Within the plot, glucose import (from extracellular to periplasm) has been indicated for reference.

**Figure S4.**
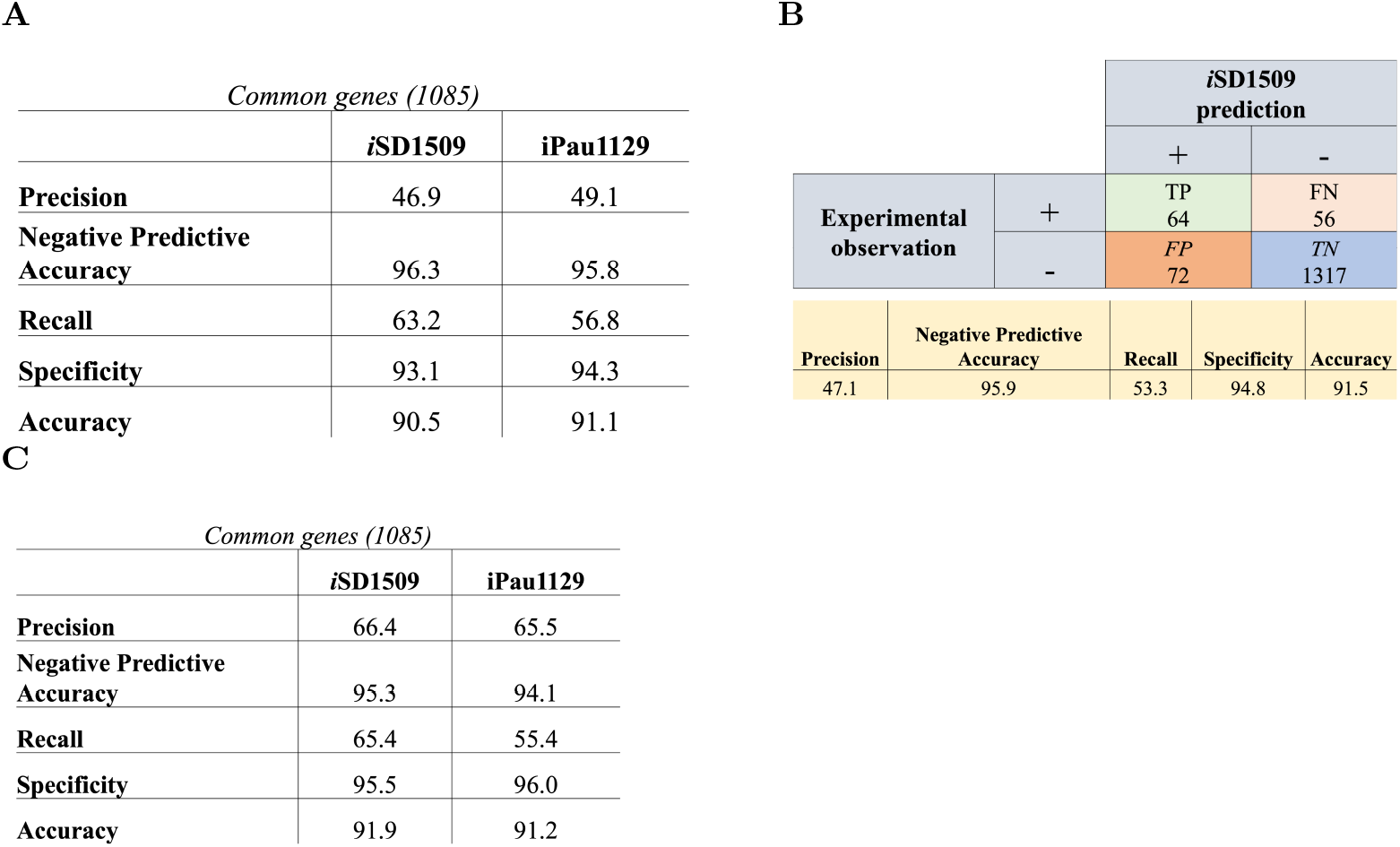
Comparison of gene essentiality between *i*SD1509 and iPau1129. (*A*) Comparing common genes between the two models for dataset retreived Liberati *et al*.[48]. (*B*) Computation of gene essentiality for all the genes present in *i*SD1509 using the same dataset [48]. (*C*) Comparing common core essential genes (from Poulsen *et al*. [35]) between the two models.

**Figure S5.**
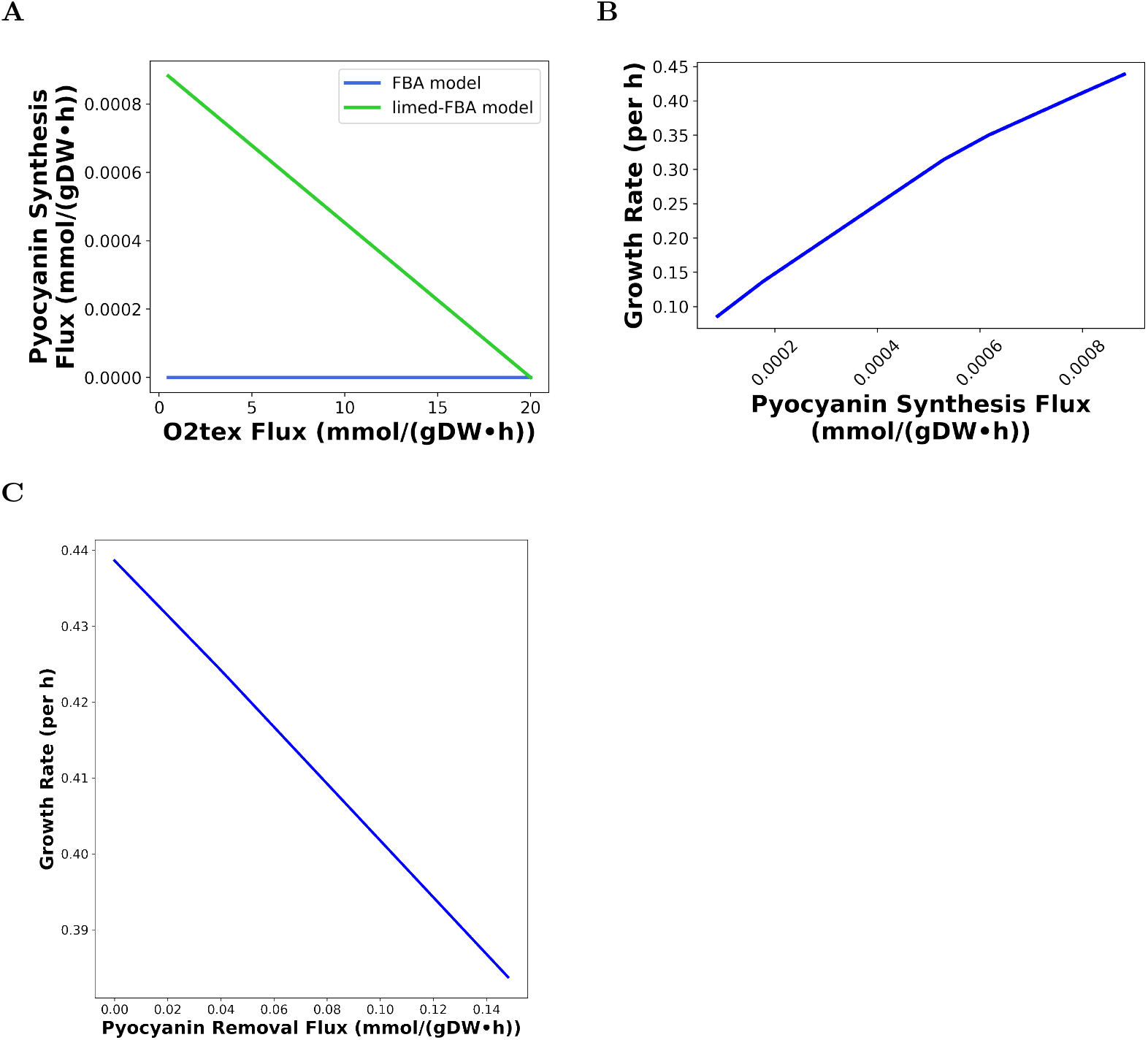
Using *i*SD1509_limed to demonstrate that the phenazine production is crucial for the pathogen survival. (*A*) Demonstration of the application of limed-FBA that predicted phenazine production is higher at lower oxygen availability condition. Normal FBA (blue) cannot predict phenazine biosynthesis as a function of oxygen availability. (*B*) In oxygen-limited condition (O2tex upper bound: 0.5 mmol gDW-1 hr-1), the biosynthesis of phenazine can affect the growth of the pathogen considerably. (*C*) A simple formulation of pyocyanin removal (i.e., a demand reaction) was added to the model, and the simulations were performed in oxygen-limited condition (O2tex upper bound: 0.5 mmol gDW-1 hr-1). As the pyocyanin removal flux increased, the model predicted that the biomass production of *P. aeruginosa* also lowered. Please note that this simulation (5*C*) can also be performed using the FBA model.

**Figure S6.**
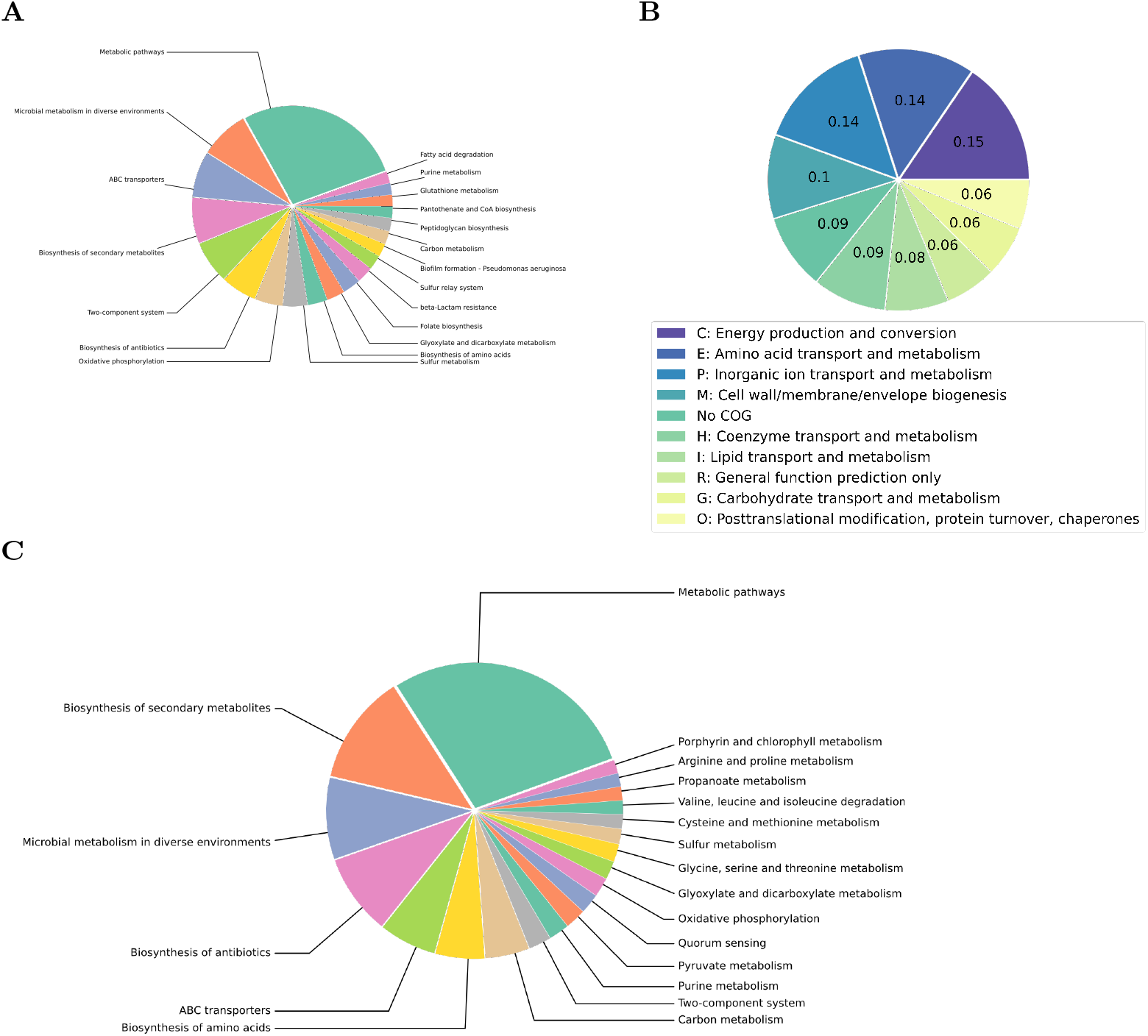
Pathway enrichment of the genes present in *i*SD1509. (*A*) The top 20 KEGG pathways of unique genes in *i*SD1509. (*B*) The top 20 COG categories of unique genes in *i*SD1509. (*C*) The top 20 KEGG pathways of all the genes present in *i*SD1509.

**Table S1.** Minimal media composition for various experiments. For model assessment and gluconate prediction study, glucose was used as the carbon substrate. For the drug potentiation/protection investigation, either fumarate or glyoxlate were used as carbon substrates.

| Model Assessment and Gluconate Prediction | Drug Potentiation/Protection |
| --- | --- |
| H<sub>2</sub>O | H<sub>2</sub>O |
| PO<sub>4</sub><sup>3-</sup> | PO<sub>4</sub><sup>3-</sup> |
| CO<sub>2</sub> | CO<sub>2</sub> |
| Fe<sup>2+</sup> | Fe<sup>2+</sup> |
| SO<sub>4</sub><sup>2-</sup> | SO<sub>4</sub><sup>2-</sup> |
| H<sup>+</sup> | H<sup>+</sup> |
| K<sup>+</sup> | K<sup>+</sup> |
| Na<sup>+</sup> | Na<sup>+</sup> |
| NH<sub>4</sub><sup>-</sup> | NH<sub>4</sub><sup>-</sup> |
| O<sub>2</sub> | O<sub>2</sub> |
| Mg<sup>2+</sup> | Mg<sup>2+</sup> |
|  | Citrate |

**Table S2.** Strains and the models used for the determination of gene-protein-reaction (GPR) associations for *i*SD1509 model. The strains are prioritized by their Average Nucleotide Identity (ANI) score (from strain 1 (*P. aeruginosa* PA14) to strain 2 (other strains)) as computed in the JGI server.

| Priority | Strain Name | Genome ID | ANI1→2 | Models used |
| --- | --- | --- | --- | --- |
| 1 | *Pseudomonas putida* KT2440 | 637000222 | 77.759 | iJN1462, iJN746 |
| 2 | *Escherichia coli* UTI89 | 637000109 | 67.768 | iUTI89_1310 |
| 3 | *Escherichia coli* str. K-12 substr. MG1655 | 646311926 | 67.712 | iML1515, iJR904, iAF1260b |
| 4 | *Escherichia coli* str. K-12 substr. W3110 | 637000110 | 67.702 | iY75_1357 |
| 5 | *Escherichia coli* ED1a | 643348545 | 67.698 | iECED1_1282 |
| 6 | *Escherichia coli* KO11FL | 2513237200 | 67.675 | iEKO11_1354 |
| 7 | *Geobacter metallireducens* GS-15 | 637000119 | 66.702 | iAF987 |
| 8 | *Mycobacterium tuberculosis* H37Rv | 637000173 | 66.467 | iEK1008, iNJ661 |
| 9 | *Synechocystis* sp. PCC 6803 | 637000315 | 62.137 | iSynCJ816 |
| 10 | *Bacillus subtilis* subtilis 168 | 646311909 | 60.516 | iYO844 |
| 11 | *Clostridioides difficile* 630 | 640069308 | 59.551 | iCN900 |
| 12 | *Clostridium ljungdahlii* DSM 13528 | 648028017 | 56.670 | iHN637 |
| 13 | *Staphylococcus aureus* subsp. aureus USA300_TCH1516 | 643692036 | 56.027 | iYS854 |

## References

[1] J. Klockgether and B. Tummler. “Recent advances in understanding Pseudomonas aeruginosa as a pathogen”. In: F1000Res 6 (2017), p. 1261. issn: 2046-1402 (Print) 2046-1402 (Linking). doi: 10.12688/f1000research.10506.1. url: https://www.ncbi.nlm.nih.gov/pubmed/28794863.

[2] Z. Pang et al. “Antibiotic resistance in Pseudomonas aeruginosa: mechanisms and alternative therapeutic strategies”. In: Biotechnol Adv 37.1 (2019), pp. 177–192. issn: 1873-1899 (Electronic) 0734-9750 (Linking). doi: 10.1016/j.biotechadv.2018.11.013. url: https://www.ncbi.nlm.nih.gov/pubmed/30500353.

[3] M. Bassetti et al. “How to manage Pseudomonas aeruginosa infections”. In: Drugs Context 7 (2018), p. 212527. issn: 1745-1981 (Print) 1740-4398 (Linking). doi: 10.7573/dic.212527. url: https://www.ncbi.nlm.nih.gov/pubmed/29872449.

[4] H. Mikkelsen, R. McMullan, and A. Filloux. “The Pseudomonas aeruginosa reference strain PA14 displays increased virulence due to a mutation in ladS”. In: PLoS One 6.12 (2011), e29113. issn: 1932-6203 (Electronic) 1932-6203 (Linking). doi: 10.1371/journal.pone.0029113. url: https://www.ncbi.nlm.nih.gov/pubmed/22216178.

[5] J. He et al. “The broad host range pathogen Pseudomonas aeruginosa strain PA14 carries two pathogenicity islands harboring plant and animal virulence genes”. In: Proc Natl Acad Sci U S A 101.8 (2004), pp. 2530–5. issn: 0027-8424 (Print) 0027-8424 (Linking). doi: 10.1073/pnas.0304622101. url: https://www.ncbi.nlm.nih.gov/pubmed/14983043.

[6] E. M. Harrison et al. “Pathogenicity islands PAPI-1 and PAPI-2 contribute individually and synergistically to the virulence of Pseudomonas aeruginosa strain PA14”. In: Infect Immun 78.4 (2010), pp. 1437–46. issn: 1098-5522 (Electronic) 0019-9567 (Linking). doi: 10.1128/IAI.00621-09. url: https://www.ncbi.nlm.nih.gov/pubmed/20123716.

[7] Bernhard Palsson. Systems biology. Cambridge university press, 2015. isbn: 1107038855.

[8] Sanjeev Dahal, Jiao Zhao, and Laurence Yang. “Genome-scale Modeling of Metabolism and Macromolecular Expression and Their Applications”. In: Biotechnology and Bioprocess Engineering (2021), pp. 1–13. issn: 1976-3816.

[9] X. Fang, C. J. Lloyd, and B. O. Palsson. “Reconstructing organisms in silico: genome-scale models and their emerging applications”. In: Nat Rev Microbiol (2020). issn: 1740-1534 (Electronic) 1740-1526 (Linking). doi: 10.1038/s41579-020-00440-4. url: https://www.ncbi.nlm.nih.gov/pubmed/32958892.

[10] L. J. Dunphy and J. A. Papin. “Biomedical applications of genome-scale metabolic network reconstructions of human pathogens”. In: Curr Opin Biotechnol 51 (2018), pp. 70–79. issn: 1879-0429 (Electronic) 0958-1669 (Linking). doi: 10.1016/j.copbio.2017.11.014. url: https://www.ncbi.nlm.nih.gov/pubmed/29223465.

[11] Mustafa Sertbas and Kutlu O Ulgen. “Genome-Scale Metabolic Modeling for Unraveling Molecular Mechanisms of High Threat Pathogens”. In: Frontiers in Cell and Developmental Biology 8 (2020).

[12] Hyun Uk Kim, Tae Yong Kim, and Sang Yup Lee. “Genome-scale metabolic network analysis and drug targeting of multi-drug resistant pathogen Acinetobacter baumannii AYE”. In: Molecular BioSystems 6.2 (2010), pp. 339–348.

[13] Jinxin Zhao et al. “Genome-Scale Metabolic Modeling Reveals Metabolic Alterations of Multidrug-Resistant Acinetobacter baumannii in a Murine Bloodstream Infection Model”. In: Microorganisms 8.11 (2020), p. 1793.

[14] Nusaibah Abdul Rahim et al. “Synergy of the Polymyxin-Chloramphenicol Combination against New Delhi Metallo-β-Lactamase-Producing Klebsiella pneumoniae Is Predominately Driven by Chloramphenicol”. In: ACS Infectious Diseases (2021).

[15] Erol S Kavvas et al. “Updated and standardized genome-scale reconstruction of Mycobacterium tuberculosis H37Rv, iEK1011, simulates flux states indicative of physiological conditions”. In: BMC systems biology 12.1 (2018), pp. 1–15.

[16] Hyun Uk Kim et al. “Integrative genome-scale metabolic analysis of Vibrio vulnificus for drug targeting and discovery”. In: Molecular systems biology 7.1 (2011), p. 460.

[17] J. A. Bartell et al. “Reconstruction of the metabolic network of Pseudomonas aeruginosa to interrogate virulence factor synthesis”. In: Nat Commun 8 (2017), p. 14631. issn: 2041-1723 (Electronic) 2041-1723 (Linking). doi: 10.1038/ncomms14631. url: https://www.ncbi.nlm.nih.gov/pubmed/28266498.

[18] Yan Zhu et al. “Genome-scale metabolic modeling of responses to polymyxins in Pseudomonas aeruginosa”. In: GigaScience 7.4 (2018), giy021.

[19] H. Arai et al. “Enzymatic characterization and in vivo function of five terminal oxidases in Pseudomonas aeruginosa”. In: J Bacteriol 196.24 (2014), pp. 4206–15. issn: 1098-5530 (Electronic) 0021-9193 (Linking). doi: 10.1128/JB.02176-14. url: https://www.ncbi.nlm.nih.gov/pubmed/25182500.

[20] T. Hirai et al. “Expression of multiple cbb3 cytochrome c oxidase isoforms by combinations of multiple isosubunits in Pseudomonas aeruginosa”. In: Proc Natl Acad Sci U S A 113.45 (2016), pp. 12815–12819. issn: 1091-6490 (Electronic) 0027-8424 (Linking). doi: 10.1073/pnas.1613308113. url: https://www.ncbi.nlm.nih.gov/pubmed/27791152.

[21] José Manuel Borrero-de Acuña et al. “Protein complex formation during denitrification by Pseudomonas aeruginosa”. In: Microbial biotechnology 10.6 (2017), pp. 1523–1534. issn: 1751-7915.

[22] J. Jo et al. “An orphan cbb3-type cytochrome oxidase subunit supports Pseudomonas aeruginosa biofilm growth and virulence”. In: Elife 6 (2017). issn: 2050-084X (Electronic) 2050-084X (Linking). doi: 10.7554/eLife.30205. url: https://www.ncbi.nlm.nih.gov/pubmed/29160206.

[23] Lars EP Dietrich et al. “Bacterial community morphogenesis is intimately linked to the intracellular redox state”. In: Journal of bacteriology 195.7 (2013), pp. 1371–1380. issn: 0021-9193.

[24] Theerthankar Das and Mike Manefield. “Pyocyanin promotes extracellular DNA release in Pseudomonas aeruginosa”. In: PloS one 7.10 (2012), e46718. issn: 1932-6203.

[25] Kyle C Costa et al. “Pyocyanin degradation by a tautomerizing demethylase inhibits Pseudomonas aeruginosa biofilms”. In: Science 355.6321 (2017), pp. 170–173. issn: 0036-8075.

[26] Charles J Norsigian et al. “BiGG Models 2020: multi-strain genome-scale models and expansion across the phylogenetic tree”. In: Nucleic acids research 48.D1 (2020), pp. D402–D406.

[27] Jonathan M Dreyfuss et al. “Reconstruction and validation of a genome-scale metabolic model for the filamentous fungus Neurospora crassa using FARM”. In: PLoS Comput Biol 9.7 (2013), e1003126. issn: 1553-7358.

[28] Ali Ebrahim et al. “COBRApy: COnstraints-based reconstruction and analysis for python”. In: BMC systems biology 7.1 (2013), p. 74. issn: 1752-0509.

[29] Nathan E Lewis et al. “Omic data from evolved E. coli are consistent with computed optimal growth from genome-scale models”. In: Molecular systems biology 6.1 (2010), p. 390. issn: 1744-4292.

[30] R Mahadevan and CH Schilling. “The effects of alternate optimal solutions in constraint-based genome-scale metabolic models”. In: Metabolic engineering 5.4 (2003), pp. 264–276. issn: 1096-7176.

[31] Volker Behrends et al. “Metabolite profiling to characterize disease-related bacteria: gluconate excretion by Pseudomonas aeruginosa mutants and clinical isolates from cystic fibrosis patients”. In: Journal of Biological Chemistry 288.21 (2013), pp. 15098–15109. issn: 0021-9258.

[32] Elad Noor et al. “Consistent estimation of Gibbs energy using component contributions”. In: PLoS Comput Biol 9.7 (2013), e1003098. issn: 1553-7358.

[33] Matthew A Oberhardt et al. “Genome-scale metabolic network analysis of the opportunistic pathogen Pseudomonas aeruginosa PAO1”. In: Journal of bacteriology 190.8 (2008), pp. 2790–2803. issn: 0021-9193.

[34] M. Kohlstedt and C. Wittmann. “GC-MS-based (13)C metabolic flux analysis resolves the parallel and cyclic glucose metabolism of Pseudomonas putida KT2440 and Pseudomonas aeruginosa PAO1”. In: Metab Eng 54 (2019), pp. 35–53. issn: 1096-7184 (Electronic) 1096-7176 (Linking). doi: 10.1016/j.ymben.2019.01.008. url: https://www.ncbi.nlm.nih.gov/pubmed/30831266.

[35] B. E. Poulsen et al. “Defining the core essential genome of Pseudomonas aeruginosa”. In: Proc Natl Acad Sci U S A 116.20 (2019), pp. 10072–10080. issn: 1091-6490 (Electronic) 0027-8424 (Linking). doi: 10.1073/pnas.1900570116. url: https://www.ncbi.nlm.nih.gov/pubmed/31036669.

[36] L. J. Dunphy, P. Yen, and J. A. Papin. “Integrated Experimental and Computational Analyses Reveal Differential Metabolic Functionality in Antibiotic-Resistant Pseudomonas aeruginosa”. In: Cell Syst 8.1 (2019), 3–14 e3. issn: 2405-4712 (Print) 2405-4712 (Linking). doi: 10.1016/j.cels.2018.12.002. url: https://www.ncbi.nlm.nih.gov/pubmed/30611675.

[37] C. D. Vo et al. “The O2-independent pathway of ubiquinone biosynthesis is essential for denitrification in Pseudomonas aeruginosa”. In: J Biol Chem 295.27 (2020), pp. 9021–9032. issn: 1083-351X (Electronic) 0021-9258 (Linking). doi: 10.1074/jbc.RA120.013748. url: https://www.ncbi.nlm.nih.gov/pubmed/32409583.

[38] S. S. Abby et al. “Advances in bacterial pathways for the biosynthesis of ubiquinone”. In: Biochim Biophys Acta Bioenerg 1861.11 (2020), p. 148259. issn: 1879-2650 (Electronic) 0005-2728 (Linking). doi: 10.1016/j.bbabio.2020.148259. url: https://www.ncbi.nlm.nih.gov/pubmed/32663475.

[39] I Koike and A Hattori. “Energy yield of denitrification: an estimate from growth yield in continuous cultures of Pseudomonas denitrificans under nitrate-, nitrite-and nitrous oxide-limited conditions”. In: Microbiology 88.1 (1975), pp. 11–19. issn: 1350-0872.

[40] Nathaniel R Glasser, Scott H Saunders, and Dianne K Newman. “The colorful world of extracellular electron shuttles”. In: Annual review of microbiology 71 (2017), pp. 731–751. issn: 0066-4227.

[41] E Tjeerd van Rij et al. “Influence of environmental conditions on the production of phenazine-1-carboxamide by Pseudomonas chlororaphis PCL1391”. In: Molecular Plant-Microbe Interactions 17.5 (2004), pp. 557–566. issn: 0894-0282.

[42] Nathaniel R Glasser et al. “The pyruvate and -ketoglutarate dehydrogenase complexes of Pseudomonas aeruginosa catalyze pyocyanin and phenazine-1-carboxylic acid reduction via the subunit dihydrolipoamide dehydrogenase”. In: Journal of Biological Chemistry 292.13 (2017), pp. 5593–5607. issn: 0021-9258.

[43] Eric E Smith et al. “Genetic adaptation by Pseudomonas aeruginosa to the airways of cystic fibrosis patients”. In: Proceedings of the National Academy of Sciences 103.22 (2006), pp. 8487–8492. issn: 0027-8424.

[44] Julie Jeukens et al. “Comparative genomics of isolates of a Pseudomonas aeruginosa epidemic strain associated with chronic lung infections of cystic fibrosis patients”. In: PloS one 9.2 (2014), e87611. issn: 1932-6203.

[45] Sylvain Meylan et al. “Carbon sources tune antibiotic susceptibility in Pseudomonas aeruginosa via tricarboxylic acid cycle control”. In: Cell chemical biology 24.2 (2017), pp. 195–206. issn: 2451-9456.

[46] J. D. Orth et al. “A comprehensive genome-scale reconstruction of Escherichia coli metabolism–2011”. In: Mol Syst Biol 7 (2011), p. 535. issn: 1744-4292 (Electronic) 1744-4292 (Linking). doi: 10.1038/msb.2011.65. url: https://www.ncbi.nlm.nih.gov/pubmed/21988831.

[47] D. Machado et al. “Fast automated reconstruction of genome-scale metabolic models for microbial species and communities”. In: Nucleic Acids Res 46.15 (2018), pp. 7542–7553. issn: 1362-4962 (Electronic) 0305-1048 (Linking). doi: 10.1093/nar/gky537. url: https://www.ncbi.nlm.nih.gov/pubmed/30192979.

[48] Nicole T Liberati et al. “An ordered, nonredundant library of Pseudomonas aeruginosa strain PA14 transposon insertion mutants”. In: Proceedings of the National Academy of Sciences 103.8 (2006), pp. 2833–2838. issn: 0027-8424.

[49] J. Nogales et al. “High-quality genome-scale metabolic modelling of Pseudomonas putida highlights its broad metabolic capabilities”. In: Environ Microbiol 22.1 (2020), pp. 255–269. issn: 1462-2920 (Electronic) 1462-2912 (Linking). doi: 10.1111/1462-2920.14843. url: https://www.ncbi.nlm.nih.gov/pubmed/31657101.

[50] Jonathan M Monk et al. “i ML1515, a knowledgebase that computes Escherichia coli traits”. In: Nature biotechnology 35.10 (2017), pp. 904–908. issn: 1546-1696.

[51] Samuel MD Seaver et al. “The ModelSEED Biochemistry Database for the integration of metabolic annotations and the reconstruction, comparison and analysis of metabolic models for plants, fungi and microbes”. In: Nucleic acids research 49.D1 (2021), pp. D575–D588. issn: 0305-1048.

[52] I-Min A Chen et al. “The IMG/M data management and analysis system v. 6.0: new tools and advanced capabilities”. In: Nucleic Acids Research 49.D1 (2021), pp. D751–D763. issn: 0305-1048.

[53] Minoru Kanehisa et al. “KEGG: integrating viruses and cellular organisms”. In: Nucleic Acids Research 49.D1 (2021), pp. D545–D551. issn: 0305-1048.

[54] Pan-Jun Kim et al. “Metabolite essentiality elucidates robustness of Escherichia coli metabolism”. In: Proceedings of the National Academy of Sciences 104.34 (2007), pp. 13638–13642. issn: 0027-8424.

[55] Jan Schellenberger and Bernhard Ø Palsson. “Use of randomized sampling for analysis of metabolic networks”. In: Journal of biological chemistry 284.9 (2009), pp. 5457–5461. issn: 0021-9258.

[56] Wout Megchelenbrink, Martijn Huynen, and Elena Marchiori. “optGpSampler: an improved tool for uniformly sampling the solution-space of genome-scale metabolic networks”. In: PloS one 9.2 (2014), e86587. issn: 1932-6203.

[57] Zachary A King et al. “Escher: a web application for building, sharing, and embedding data-rich visualizations of biological pathways”. In: PLoS Comput Biol 11.8 (2015), e1004321. issn: 1553-7358.

